# Stress-induced switch in small extracellular vesicle secretion: from constitutive “torn bag mechanism” to exocytosis

**DOI:** 10.1101/2025.10.29.685290

**Authors:** Dorina Lenzinger, Lilla Lankovics, István Dudás, Tünde Bárkai, Zsófia Szász, Krisztina V Vukman, Kelsey Fletcher, Attila Csomos, Zoltán Mucsi, Edina Bugyik, Csaba Cserép, Ádám Dénes, Szilvia Bősze, Edit I Buzás, Tamás Visnovitz

## Abstract

The biogenesis of small extracellular vesicles (sEVs) is only partially understood. Our recent findings provide evidence that a newly described sEV secretion pathway, the amphiectosome release and the “torn bag mechanism”, is present in all tested cell lines and in mouse liver and kidney. Surprisingly, in *in situ* fixed steady-state cells, transmission electron microscopy did not reveal the classical exosome secretion route, the sEV release via exocytosis of multivesicular endosomes (MVEs). In the current study, we investigated which parameters influence the activation of the two distinct sEV release mechanisms. Our results show that under stress conditions (such as Ca²⁺ ionophore-induced membrane stress or metabolic stress-induced by serum starvation), exocytosis of MVEs is activated, while this process is absent in steady-state conditions. By silencing ATG5 (a key regulator of autophagy) and RAB27a (essential small GTPase for MVE exocytosis), we selectively modulated these two mechanisms. Amphiectosome release depended on both autophagy and ATG5, while exocytosis of MVE was autophagy-independent but RAB27a-dependent. Our findings suggest that sEV release via the “torn bag mechanism” is a general and essential secretion pathway in non-stressed, steady-state mammalian cells, while stress conditions induce the sEV release via MVE exocytosis.

## Introduction

Extracellular vesicles (EVs) are particles enclosed by a phospholipid bilayer that facilitate intercellular communication [1, 2], and exosomes are a subset of EVs of endosomal origin [1, 3]. These small EVs (sEVs), typically less than 200 nm in diameter, are formed by the inward budding of the limiting membrane of late endosomes or multivesicular bodies (MVBs) [4, 5]. Exosome release is considered to be associated with the exocytosis of intraluminal vesicles (ILVs) of multivesicular endosomes (MVEs) [4, 6]. In contrast, ectosomes bud directly from the cell surface surrounded by the plasma membrane-derived phospholipid membrane [1, 3, 6]. Recent studies have uncovered an unexpected complexity in EV biogenesis [3]. We recently reported that the endosomal and plasma membrane origin of EVs are not necessarily mutually exclusive [7, 8]. In colorectal carcinoma cells, we described the ectocytotic release of “MVB-like sEV clusters” containing ILVs of endosomal origin. [7].

Our latest findings also provided evidence for a novel, active, autophagy-dependent, exocytosis-independent sEV secretion pathway, termed amphiectosome release and “torn bag mechanism” [8]. Interestingly, both the ILVs of amphiectosomes and the sEVs separated from conditioned media were exclusively positive either for the classical sEV markers (such as CD63, CD81, ALIX and TSG101) or LC3B, suggesting an MVB or autophagy origin of these vesicles. According to our proposed model, upon fusion of MVBs with autophagosomes, fragments of the autophagosomal inner membrane curl up to form LC3B-positive ILVs within amphisomes. In contrast, the CD63-positive sEVs are of presumed MVB origin. The amphisomes are shed from the plasma membrane via ectocytosis as amphiectosomes, which subsequently release their ILVs into the extracellular space through the “torn bag mechanism” [8]. The amphiectosome release and “torn bag” EV secretion mechanism appear to be frequent biological processes as we identified them in all tested cell cultures as well as in mouse kidney and liver tissues [8]. Although the classical exosome release mechanism is linked to the exocytosis of MVBs [4, 6] or amphisomes [6, 9], there is relatively limited ultrastructural evidence supporting this process [10–14]. Most of these studies have been conducted on special cell types such as reticulocytes [10, 11], immune cells [12], platelets [14] or tumour cells [13].

In contrast to the data in the literature, we have not detected obvious MVE exocytosis in cell cultures or in ultrathin sections of various mammalian tissues under steady-state conditions. Recent findings suggest that Ca²⁺ influx can trigger MVB exocytosis and exosome release, potentially playing a role in plasma membrane repair through the ANX6 protein [15]. Here we investigated the “torn bag mechanism” and MVE exocytosis in sEV secretion. Specifically, we examined which parameters influence the activation of these two distinct sEV secretion pathways and found that exocytosis is induced by cellular stress.

## Materials and Methods

### Cell lines

The HEK293 human embryonic kidney cell line (#85120602, Batch No: 18E026) was purchased from the European Collection of Authenticated Cell Cultures (ECACC) through their distributor (Sigma), the HEK293T-PalmGFP human embryonic kidney cells were kindly provided by Charles P. Lai [16]. The HEK293T-PalmGFP-LC3RFP cell line was developed by our research group previously [8]. All HEK293 cell lines were grown in DMEM (Gibco) with 10% fetal bovine serum (FBS, BioSera) in the presence of 2 µmol/mL l-glutamine (Sigma), 100 U/mL of Penicillin and 100 µg/mL Streptomycin (Sigma) [17]. Before microscopy sample preparation the cells were cultured on the surface of 0.02% gelatin (Sigma) and 5 μg/mL fibronectin (Invitrogen) coated glass coverslips (VWR) [18]. In order to minimize the genetic drift, master cell banks (MCB) and subsequently, working cell banks (WCBs) were established. Vials of the characterised WCBs were used for experiments. The cell banks were tested for Mycoplasma infection by PCR [18].

### Mice

C57BL/6 mice were purchased from The Jackson Laboratory and bred in a pathogen-free animal facility at the Institute of Genetics, Cell and Immunobiology, Semmelweis University. For transmission electron microscopy, 12-week-old male mice were used and sacrificed in accordance with the Animal Experimentation Code of Semmelweis University.

### Transmission electron microscopy

Tissue samples from mouse organs (approximately 1 mm³) were immersed and fixed in 4% glutaraldehyde (Sigma) for at least 24 hours at 4 °C, followed by post-fixation in 1% osmium tetroxide (SPI) for 2 hours at room temperature (RT). For embedding, an EPON resin mixture (SPI) was applied as described previously [8, 19]. Sample preparation of HEK293 cells was performed on the surface of glass coverslips placed in a 24-well untreated cell culture plate (Eppendorf). For monolayer cell cultures, propylene oxide (typically used as a solvent for EPON resin) was omitted and replaced with absolute ethanol. After polymerization of the resin (at least 48 hours at 56 °C), the glass coverslips were removed using liquid nitrogen. Ultrathin sections (50-70 nm) were made using an RMC Boeckeler PowerTome ultramicrotome. Sections were post-contrasted with UranyLess (Electron Microscopy Sciences) for 10 minutes at RT, followed by 2 minutes in lead citrate (RT) [20]. Images were captured using a JEM-1010 transmission electron microscope (JEOL) equipped with a 3-megapixel Veleta digital camera with the help of Olympus iTEM software (version 5.1).

### Immunofluorescent microscopies

Confocal microscopy was performed as previously described [8]. Cells were cultured on a gelatin/fibronectin-coated surface. To study the cellular microenvironment around the cultured cells, the cultures were not washed before fixation; both cells and their microenvironment were fixed *“in situ”*. Without changing the culture medium, 8% paraformaldehyde (PFA) in PBS was added in an equal volume to achieve a final concentration of 4%. After fixation (20 min, RT), samples were washed twice with PBS (5 min each, RT) and once with 50 mM glycine in PBS (5 min, RT). Blocking and permeabilization were carried out using 10% FBS with 0.1% Triton X-100 (Sigma) in PBS for 1h at RT. Primary antibodies were applied at 1:200 dilution in the same blocking/permeabilization solution and incubated overnight at 4 °C. Excess primary antibodies were removed by two washes (5 min each) with the blocking/permeabilization solution at RT, followed by one wash (5 min, RT) with 1% FBS in PBS. Secondary antibodies were applied at 1:500-1:1000 dilution in 1% FBS in PBS for 1h at RT. Unbound secondary antibodies were removed by two washes (5 min each) with PBS and two washes (5 min each) with distilled water. Details of the antibodies used are provided in Table 1. Samples were mounted in SlowFade™ Diamond with DAPI (Invitrogen) or ProLong™ Diamond with DAPI (Invitrogen). For quantification of the cellular microenvironment, images were acquired using an Olympus FluoView FV4000 inverted confocal microscope (EVIDENT) equipped with a 60x oil immersion objective (UPLXAPO60XO, NA 1.42, WD 0.15 mm). Samples were sequentially illuminated with 405 nm, 488 nm, 561 nm and 647 nm lasers. Images were scanned using galvanometer scanners at 2048 × 2048 pixel resolution. Signals were detected with broadband and red-shifted SilVIR detectors at a dwell time of 2 μs/pixel. Image analysis was performed using Fiji software.

**Table 1.** List of antibodies used in the study.

| **Antibody** | **Manufacturer** | **Identifiers** | **Dilutions** |
| --- | --- | --- | --- |
| monoclonal anti-LC3B (rabbit) | Sigma/Merck | ZRB100<br>clone: 12K5 | WB: 1:1000<br>IF: 1:200<br>STED: 1:200 |
| monoclonal anti-CD63 (mouse) | Santa Cruz Biotechnology | MX-49.129.5<br>clone: sc-5275<br>RRID:AB_627877 | IF: 1:200<br>STED: 1:200 |
| polyclonal anti-CD63 (C terminal) (rabbit) | Sigma/Merck | SAB2109138 | WB: 1:500 |
| polyclonal anti-γ-actin (N terminal) (rabbit) | Sigma | A2103<br>RRID:AB_476694 | WB: 1:1000 |
| goat anti-rabbit Star 580 | Abberior | ST580-1002-500UG<br>RRID:AB_2910107 | STED: 1:500 |
| goat anti-mouse Star 635P | Abberior | ST635P-1001-500UG<br>RRID:AB_2893232 | STED: 1:500 |
| monoclonal anti-ATG5 (rabbit) | Sigma/Merck | ZRB1596<br>clone 1L18 | WB: 1:1000 |
| polyclonal anti-RAB27a (rabbit) | Invitrogen | PA5-79904<br>RRID:AB_2747019 | WB: 1:1000 |
| polyclonal goat anti-rabbit IgG Fc (HRP) | Abcam | ab97200<br>RRID:AB_10679899 | WB: 1:10000 |

For multi-channel stimulated emission depletion (STED) nanoscopy SlowFade™ Diamond Antifade Mountant (Thermo Fisher Scientific) was applied. Imaging was carried out on an Abberior Instruments Facility Line STED microscope system built on an Olympus IX83 fully motorized inverted microscope base. The system was equipped with a ZDC-830 TrueFocus Z-drift compensator, an IX3-SSU ultrasonic stage, a QUADScan beam scanner, avalanche photodiode (APD) detectors and a UPLXAPO60XO 60x oil immersion objective (NA 1.42). Excitation was performed using 488 nm, 561 nm and 640 nm solid-state lasers, while a 775 nm solid-state laser was used for STED depletion. Image acquisition was controlled using the Imspector data acquisition software (v.16.3.14278-w2129-win64). All other slides were examined using a Leica SP8 Lightning confocal microscope in adaptive lightning mode with a HC PL APO CS2 63x/1.40 oil immersion objective and hybrid detector. Lookup tables (LUTs) were kept linear throughout the study. Image analysis was performed using Leica LAS X software.

### Modulation of different sEV release pathways

Exocytosis of MVEs was induced using the Ca²⁺ ionophore A23187 (Sigma) [15]. A 1 mM stock solution was prepared in dimethyl sulfoxide (DMSO, Serva). Working solutions at final concentrations of 100 nM, 250 nM, 500 nM and 1000 nM were made by diluting the stock in fresh, pre-warmed, serum- and antibiotic-free cell culture medium. Cells were treated for 2.5, 5 and 10 min, after which both the cells and their microenvironment were fixed with 4% PFA. For nanoparticle tracking analysis (NTA), high-resolution flow cytometry and Western blotting, conditioned media were collected. The collected media were centrifuged at 300g for 5 min at RT to remove cellular contaminants and then stored on ice until further processing. Autophagy was induced by chloroquine as described previously [8].

### Viability measurements

We used the resazurin assay to evaluate the effect of the Ca²⁺ ionophore on cell viability through mitochondrial metabolic activity [8, 18]. Equal numbers of cells were plated in a 48-well plate (Eppendorf). At 80% confluence, the culture medium was replaced with medium containing the Ca²⁺ ionophore. After the indicated incubation times, the medium was replaced with fresh medium supplemented with serum, antibiotics and 10 µg resazurin (Sigma). Following 3h of incubation, the cells were allowed to recover, after which fluorescence changes were measured by Fluoroskan FL fluorometer (Thermo Scientific). To assess the immediate effect of Ca²⁺ ionophore treatment on cell viability, we performed a CellTiter-Glo assay (Promega), which measures cellular ATP levels. Equal numbers of cells were plated into 96-well plate (TPP). Upon reaching 80% confluence, the culture medium was replaced and Ca²⁺ ionophore treatments were applied. At the specified time points, the CellTiter-Glo assay was performed according to the manufacturer’s instructions. The final products were transferred to a white plate to minimize background noise. Luminescence was measured using Fluoroskan FL fluorometer (Thermo Scientific) [21].

### Nanoparticle Tracking Analysis (NTA)

Particle concentration and size distribution were determined using a ZetaView PMX120 nanoparticle tracking analysis (NTA) instrument (Particle Metrix) in both scatter and fluorescence modes. Lipophilic EVs were distinguished from non-EV particles using BioxMLYellow (Bioxol), a self-quenchable fluorescent dye [22]. For fluorescence measurements, samples were labelled with 5 nM dye for approximately 5 min. Instrument settings are provided in Supplementary Table 1.

### High resolution flow cytometry

EV secretion from A23187-induced cells was monitored using high-resolution flow cytometry (Apogee Micro-PLUS, Apogee Flow Systems). EVs were identified based on the presence of phospholipid membranes, using BioxMLRed (Bioxol), a lipophilic and self-quenching fluorescent dye, and by their sensitivity to Triton X-100^®^ (Molar Chemicals), as previously described [18, 23]. A detailed description of the high-resolution flow cytometry settings is provided in Supplementary Table 2. Data analysis was performed as shown in Supplementary Figure 2, using FlowJo_V10 software.

### Holographic Microscopy

Morphometric changes induced by Ca²⁺ ionophore treatment were analysed using a holographic transmission microscope (HoloMonitor M4; Phase Holographic Imaging AB) [21, 24]. Changes in cellular morphology, including average cell area, shape irregularity, texture contrast, optical thickness and optical volume were monitored and quantified. HEK293 cells were cultured and analysed at approximately 60% confluence. Before imaging, the culture medium was replaced with 3 mL of serum-free medium, either containing 500 nM Ca²⁺ ionophore or without ionophore (control). Time-lapse images were acquired every 10 sec for 10 min. For image analysis, at least 70 cells per field of view were identified using the minimum error histogram-based thresholding algorithm implemented in the built-in software (HStudio M4).

### Gene silencing

ATG5 and RAB27a genes were silenced in HEK293 and HEK293T-PalmGFP-LC3RFP cells using pre-designed Silencer^®^ Select siRNAs (Ambion). Transfections were performed using Lipofectamine™ RNAiMAX (Invitrogen) according to the manufacturer’s protocol with minor modifications. Cells were maintained in antibiotic-free DMEM until they reached 30-50% confluence. RNAi duplex-Lipofectamine™ RNAiMAX complexes were prepared in serum- and antibiotic-free Opti-MEM medium (Gibco) at a final siRNA concentration of 10 nM. After 4-6 hours of transfection, 10% serum was added to the medium, and cells were incubated for an additional 3 days. Subsequently, the medium was replaced with fresh siRNA-containing Opti-MEM, and serum was reintroduced after 4-6 hours. Cells were cultured for one more day before being used in experiments. The sequences of the siRNAs are provided in Table 2.

**Table 2.**
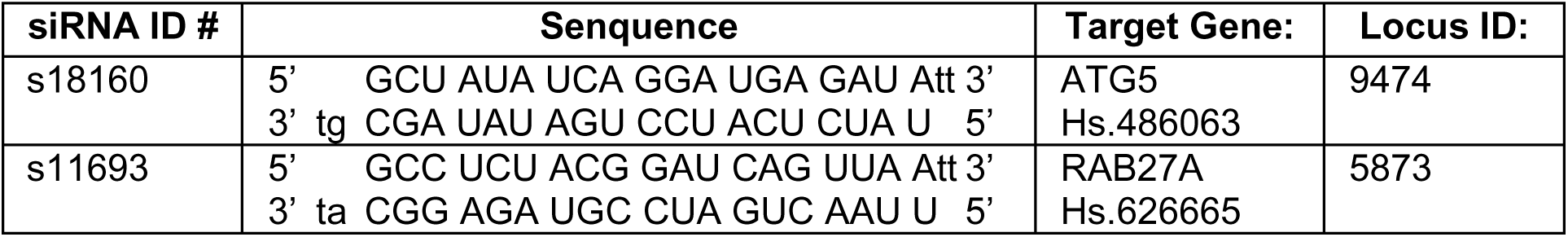
Nucleotide sequences of applied siRNAs.

### Quantitative Real-Time PCR

To assess gene silencing at the RNA level, quantitative real-time PCR (qPCR) was performed. PCR primers were designed using Primer3web (https://primer3.ut.ee/) and validated on a Kyratec SC300T PCR system (Kyratec) followed by agarose gel electrophoresis. Sequences of the PCR primers are listed in Table 3. Total RNA was extracted from HEK293 cells using the NucleoSpin® RNA Plus Kit (MACHEREY-NAGEL) according to the manufacturer’s instructions. Cells were cultured in 6-well plates (VWR) or T12.5 flasks (VWR). After mechanical detachment, cells were collected by centrifugation at 300g for 5 min at RT. Pellets were lysed in the provided lysis buffer and RNA was eluted in 30 µL of RNase-free water. RNA content and purity were determined using a NanoDrop-1000 spectrophotometer (Thermo Fisher Scientific). The reverse transcription was performed using 1 µg of total RNA and an Oligo(dT)₁₈ primer with the RevertAid First Strand cDNA Synthesis Kit (Thermo Fisher Scientific), following the manufacturer’s protocol. The qPCR was carried out using 20 ng of cDNA, 3-5 pmol of each primer and SensiFAST™ SYBR Hi-ROX Mix (BioLine) on a CFX96 Touch™ Real-Time PCR System (Bio-Rad). Data were analysed using CFX Maestro software (Bio-Rad). The amplification protocol is summarized in Table 4. Gene expression levels were normalized to the housekeeping gene GAPDH.

**Table 3.**
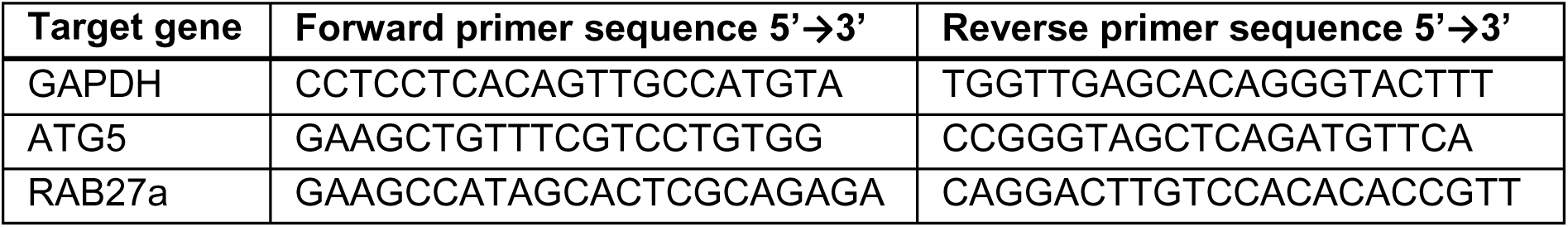
List of PCR primers:

**Table 4.**
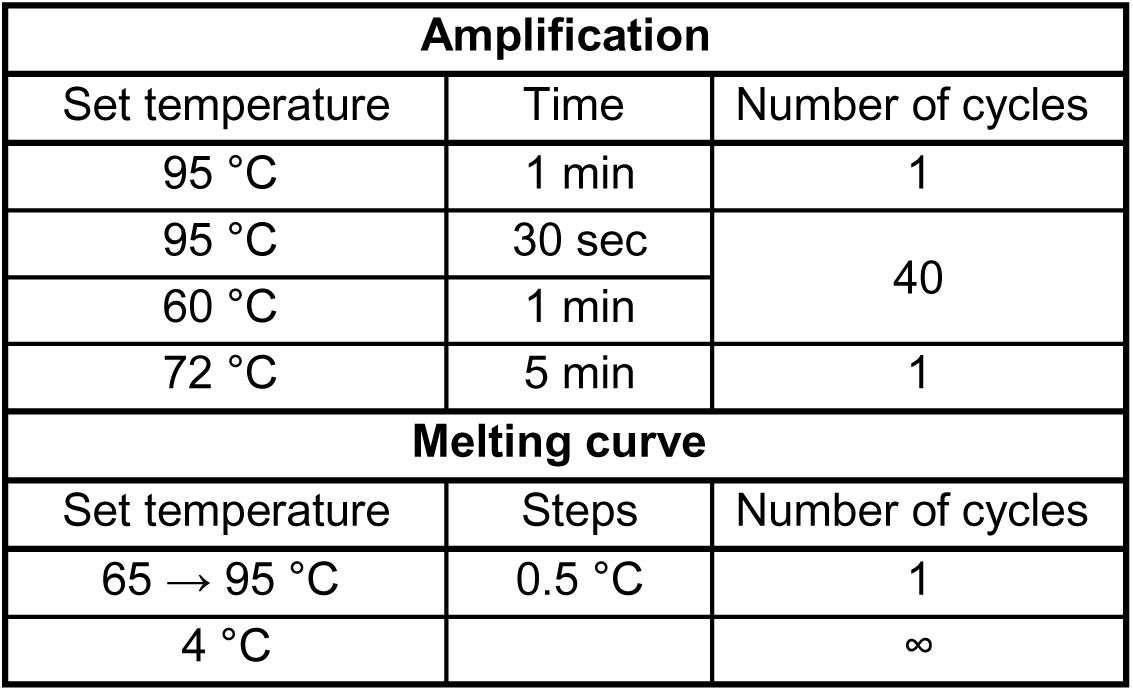
Thermal profile of qPCR reactions.

| **Amplification** |  |  |
| --- | --- | --- |
| Set temperature | Time | Number of cycles |
| 95 °C | 1 min | 1 |
| 95 °C | 30 sec | 40 |
| 60 °C | 1 min |  |
| 72 °C | 5 min | 1 |
| **Melting curve** |  |  |
| Set temperature | Steps | Number of cycles |
| 65 → 95 °C | 0.5 °C | 1 |
| 4 °C |  | ∞ |

### Western blotting

Western blotting was performed as described previously [8, 18]. Whole-cell lysates were analysed to assess silencing efficiency at the protein level. Cells were cultured in T25 flasks (Eppendorf) until reaching approximately 80% confluence. After mechanical detachment, cells were harvested by centrifugation at 300g for 5 min at RT. Pellets were lysed in radio-immunoprecipitation assay (RIPA) buffer supplemented with 1 mM PMSF (Serva) and 2 mM sodium orthovanadate (Sigma). For analysis of the extracellular environment, total protein from conditioned, serum- and antibiotic-free medium was precipitated using trichloroacetic acid (TCA) as described previously [18, 25] with minor modifications. After removing cells by centrifugation (300g, 10 min, RT), proteins were precipitated on ice for at least 1h. Precipitated proteins were collected by centrifugation at 20.000g for 20 min at 4 °C, then washed twice with -20 °C acetone. Protein pellets were resuspended in RIPA buffer as described above. Protein concentrations were determined using the Micro BCA Protein Assay Kit (Thermo Fisher Scientific) and a MULTISKAN SkyHigh Microplate Spectrophotometer (Thermo Fisher Scientific).

Proteins were separated on 12% polyacrylamide gels (acrylamide/bis-acrylamide ratio 37.5:1) using a Mini-PROTEAN system (Bio-Rad). For improved solubilisation of membrane proteins, equal volumes of 0.1% Triton X-100, Laemmli buffer, and samples were mixed as described previously [26]. Within each biological replicate, equal amounts of protein (∼15-20 µg) were loaded per lane. After electrophoresis, proteins were transferred to PVDF membranes (Serva). Membranes were blocked with 5% BSA in TBS-T for 1h at RT. Primary antibodies and their dilutions are listed in Table 1. Peroxidase-conjugated secondary antibodies were applied at 1:10.000 dilution. Signals were detected using ECL Western Blotting Substrate (Thermo Scientific) and visualized with an Imager CHEMI Premium system (VWR).

### Software and statistical analysis

For image capturing and analysis CellSense FV software (EVIDENT), LASX (Leica), Imspector (v.16.3.14278-w2129-win64, Abberior Instruments), Olympus iTEM, HStudio M4 and Image J software were used. Figures and graphs were generated using GraphPad Prism 9.4.1 and Biorender (BioRender.com). For statistical analysis, standard deviation was calculated. Unpaired two tailed Student’s t-tests and one-way ANOVA were used (* p<0.05, ** p<0.01, *** p<0.001, **** p<0.0001).

## Results

### Identification of MV-lEVs in different mouse tissues

In our very recent study, we demonstrated the presence of large multivesicular extracellular vesicles (MV-lEVs) in various cell lines as well as in mouse kidney and liver. In many cases, we confirmed that these MV-lEVs represent a novel EV subtype, with an amphisomal origin, and we termed them amphiectosomes [8]. First, here we addressed the question whether MV-lEV secretion is a general process, which can be observed in different organs and tissues of a complex organism such as the mouse. Ultrathin sections of mouse spleen (Figure 1A-C), lungs (Figure 1D-E), skeletal muscle (Figure 1F,G), heart (Figure 1H,I), small intestine (Figure 1J-L), kidney (Figure 1M) and liver (Figure 1N,O) were examined by transmission electron microscopy. MV-lEVs were detected in all analysed tissues. Furthermore, different stages of MV-lEV secretion were observed, including budding (Figure 1A, E, H, O), intact MV-lEVs in the extracellular space (Figure 1B, D, F, G, I, K, N) and release of intraluminal vesicles (Figure 1C, J, M). Thus, the characteristic “torn bag mechanism” was clearly visible in all the tested tissues and organs. In agreement with our previous findings, under steady-state conditions, secretion of MVEs by exocytosis was not detected. In the small intestine, however, secretion of sEVs via ectocytosis was observed (Figure 1L), where sEVs were released from microvilli.

**Figure 1.**
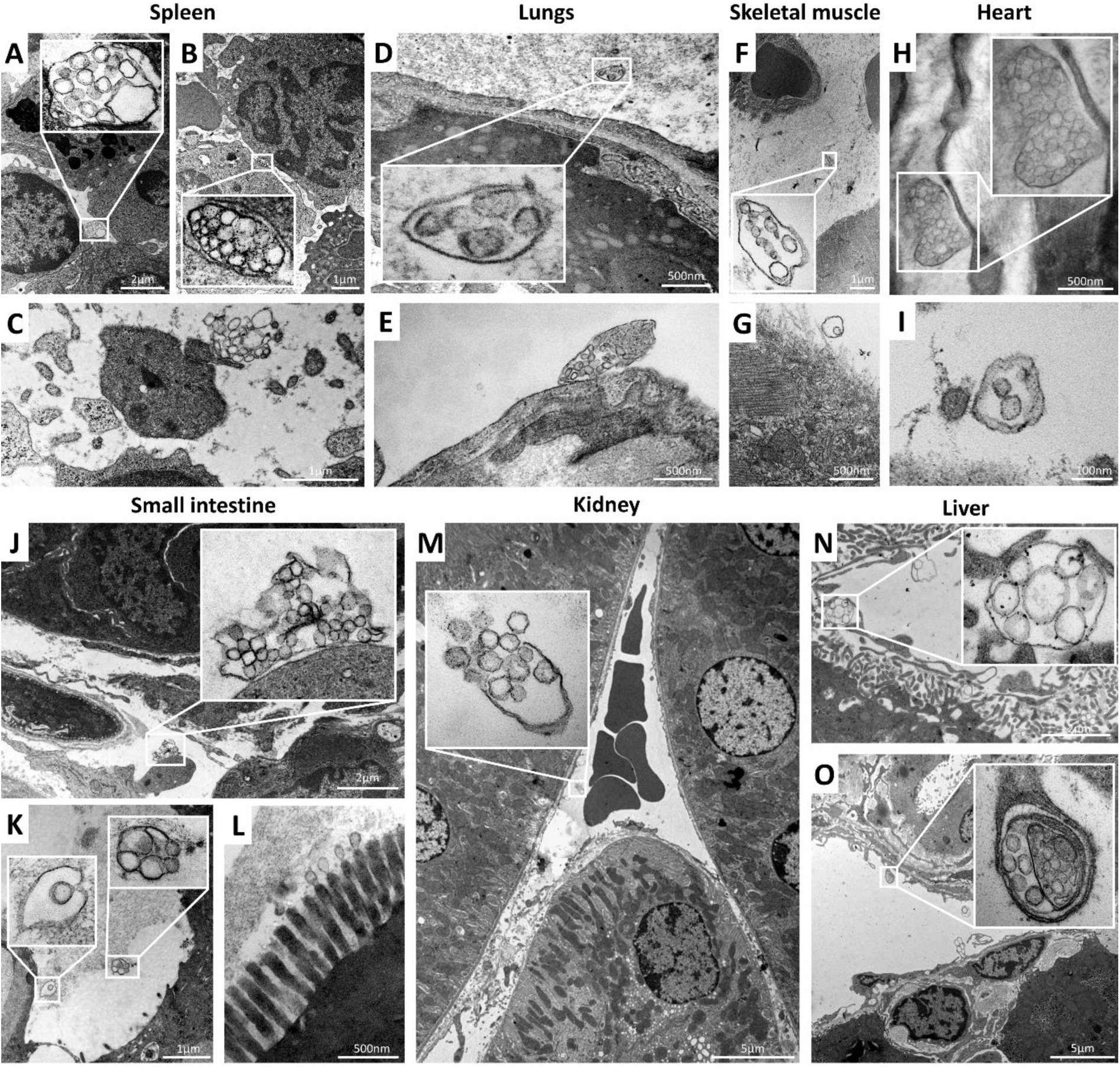
MV-lEVs in mouse organs and tissues. Transmission electron microscopy revealed MV-lEVs in spleen (A-C), lungs (D,E), skeletal muscle (F,G), heart (H,I), small intestine (J-L), kidney (M) and liver (N,O). Stages of MV-lEV secretion were observed, such as budding (A,E,H,O), intact MV-lEVs in the extracellular environment (B,D,F,G,I,K,N), and release of intraluminal vesicles (C,J,M). Under steady-state conditions, secretion of MVEs by exocytosis was not detected, while sEV secretion via ectocytosis occurred in the small intestine (L).

### Effect of Ca²⁺ ionophore on EV secretion

To investigate the effect of Ca²⁺ ionophore on EV secretion, HEK293 cells were exposed to increasing concentrations of A23187 and analysed by TEM, NTA and high-resolution flow cytometry (Figure 2). Under steady-state control conditions and at A23187 concentrations below 250 nM, the predominant sEV release pathway was the “torn bag mechanism,” whereas MVE exocytosis was not clearly detected. TEM revealed distinct stages of MV-lEV secretion, including plasma membrane budding, intact MV-lEVs in the extracellular space, release of intraluminal vesicles and empty limiting membrane balloons (Figure 2A-L). These features were consistently observed across multiple fields, confirming that MV-lEV secretion is an active and dynamic process under these conditions.

**Figure 2.**
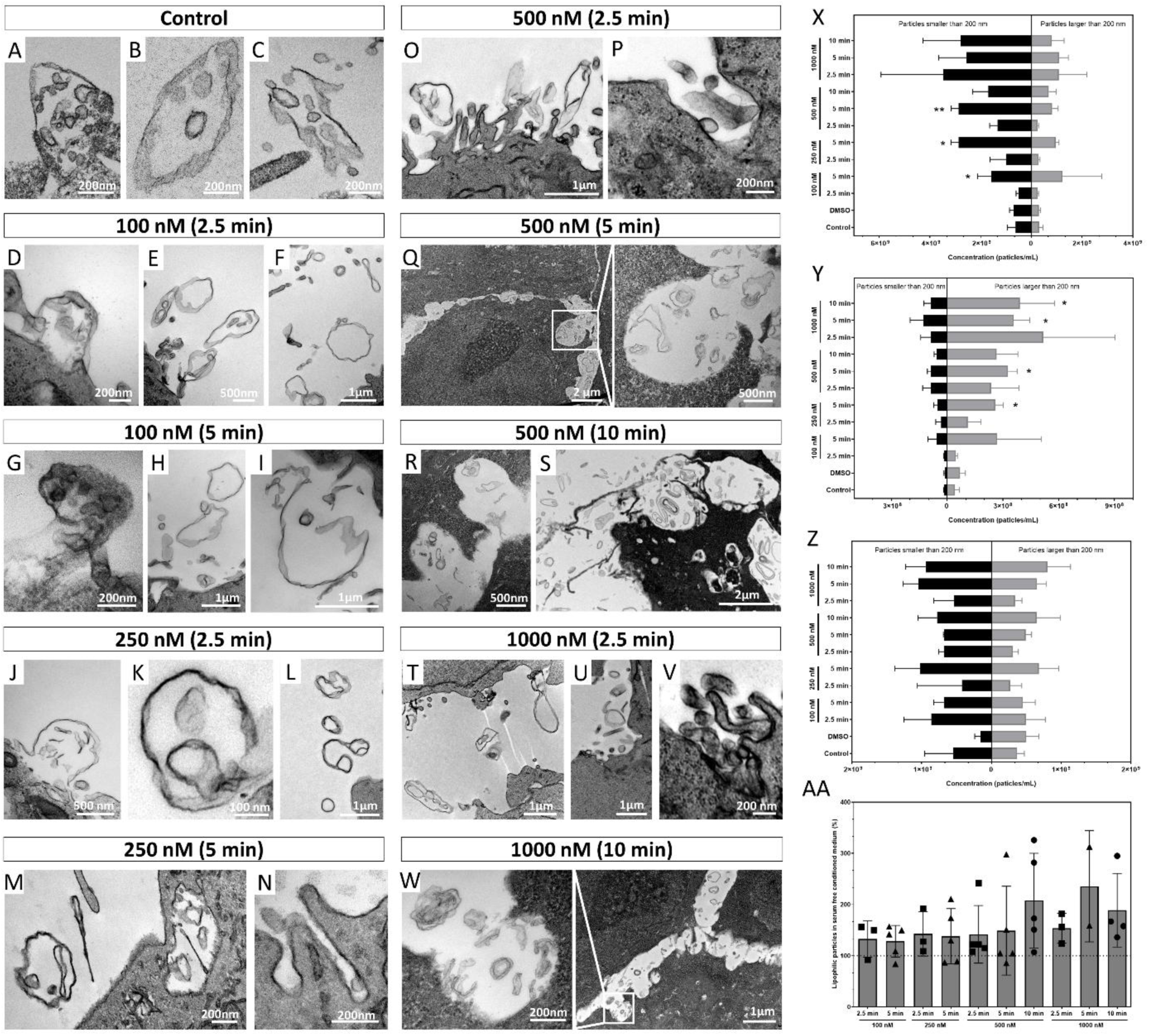
Effect of Ca²⁺ ionophore on EV secretion. The impact of A23187 Ca²⁺ ionophore treatment on HEK293 cells was assessed using transmission electron microscopy (A-W), nanoparticle tracking analysis (NTA; X-Z), and high-resolution flow cytometry (AA). Under control conditions (A-C) and at A23187 concentrations up to 250 nM (D-N), the predominant sEV secretion pathway was the “torn bag mechanism”. Multivesicular endosome (MVE) exocytosis was observed only after 5 min of treatment with 250 nM A23187 (N). Stages of MV-lEV secretion included budding (A, D, G, J), intact MV-lEVs in the extracellular space (B, E, K, M), release of intraluminal vesicles (C, I) and empty limiting membrane balloons of MV-lEVs (F, E, H, L) were recorded. NTA was applied to quantify particle release in scatter mode for sEV (X) and lEV (Y) settings and in fluorescence mode (Z). In scatter mode, both small (<200 nm; X) and larger (>200 nm; Y) particle numbers increased significantly in a concentration- and time-dependent manner up to 1000 nM A23187 concentration. No further increase was detected above 1000 nM A23187. When a lipophilic fluorescent dye (BioxML-Yellow) was applied, this significant increase was not observed in fluorescence-mode NTA (Z). Findings further validated by high-resolution flow cytometry (AA). Bar charts represent mean ± SD of 2-5 independent experiments. Statistical significance was determined using Student’s t-test and one-way ANOVA (*p < 0.05, **p < 0.01).

Importantly, at 250 nM A23187 for 5 min, we detected MVE exocytosis (Figure 2N) in addition to MV-lEV secretion and the “torn bag mechanism” (Figure 2M). With further increase in Ca²⁺ ionophore concentration and treatment duration, MVE exocytosis became the dominant sEV release mechanism (Figure 2O-W). Of note, at higher A23187 concentrations, both MV-lEV secretion (Figure 2O,T) and MVE exocytosis (Figure 2P,U,V) were observed at early time points (2.5 min), whereas at later stages (5-10 min), only MVE exocytosis and large empty membrane balloons were detected (Figure 2Q-S,W). These empty structures may originate from the limiting membranes of ruptured MV-lEVs.

For quantitative analysis of Ca²⁺ ionophore-dependent particle release, NTA and high-resolution flow cytometry were performed testing cell free supernatant, diluted in PBS. NTA in scatter mode revealed a significant, concentration- and time-dependent increase in both small (<200 nm, Figure 2X) and larger (>200 nm, Figure 2Y) particles up to 1000 nM A23187. At 1000 nM A23187 concentration, no further increase was observed in particle counts, suggesting a saturation effect. Interestingly, when lipophilic fluorescent labelling was applied, no significant increase in fluorescence was detected, although a slight trend was observed (Figure 2Z). The discrepancy between scatter and fluorescence modes suggests that a substantial fraction of released particles may lack a limiting membrane. Median (X50) size of particles in supernatant are indicated in Supplementary Figure 1.

These findings were further supported by high-resolution flow cytometry. Although an increasing trend in EV release was observed, it did not reach statistical significance (Figure 2AA). Overall, Ca²⁺ ionophore treatment significantly enhances the release of extracellular particles in a dose-dependent manner; however, only a subset of these particles appears to be real EVs. Gating strategy of EVs detection is found in Supplementary Figure 2.

### Characterisation of Ca²⁺ ionophore-induced EV secretion

To investigate the nature of the EVs released by the effects of Ca^2+^ ionophore, we tested HEK293T-PalmGFP cells. We employed a combination of immunocytochemistry, confocal microscopy, STED super-resolution microscopy and holographic microscopy. EV subtypes were identified and quantified based on PalmGFP, CD63 and LC3B markers.

Representative confocal images, which were used for quantification, are shown in Figure 3 A-D. The results reveals distinct changes in EV subpopulations following Ca^2+^ ionophore (A23187) treatment. PalmGFP fluorescence, indicating plasma membrane-derived EVs, fluctuated around control levels with minor but significant changes (Figure 3E). In contrast, CD63-positive EVs exhibited pronounced time- and concentration-dependent responses. At 2.5 min, CD63 signal continuously increased and reached saturation at 500 nM. Similar a trend was recognised at 5 min time points. Following the effect of concentration dependency, at lower concentrations (100-250 nM) CD63 fluorescence continued to increase over time, whereas at higher concentrations (500-1000 nM), a significant decrease occurred at 10 min time points (Figure 3F). The LC3B-positive EVs followed a pattern similar to PalmGFP, with an extra peak at 250 nM and 5 min. A slight and significant concentration-dependent increase was observed at 2.5 min, which plateaued at a 500 nM Ca^2+^ ionophore concentration. At higher concentrations of A23187 (500-1000 nM), a decreasing time-dependent trend was found (Figure 3G).

**Figure 3.**
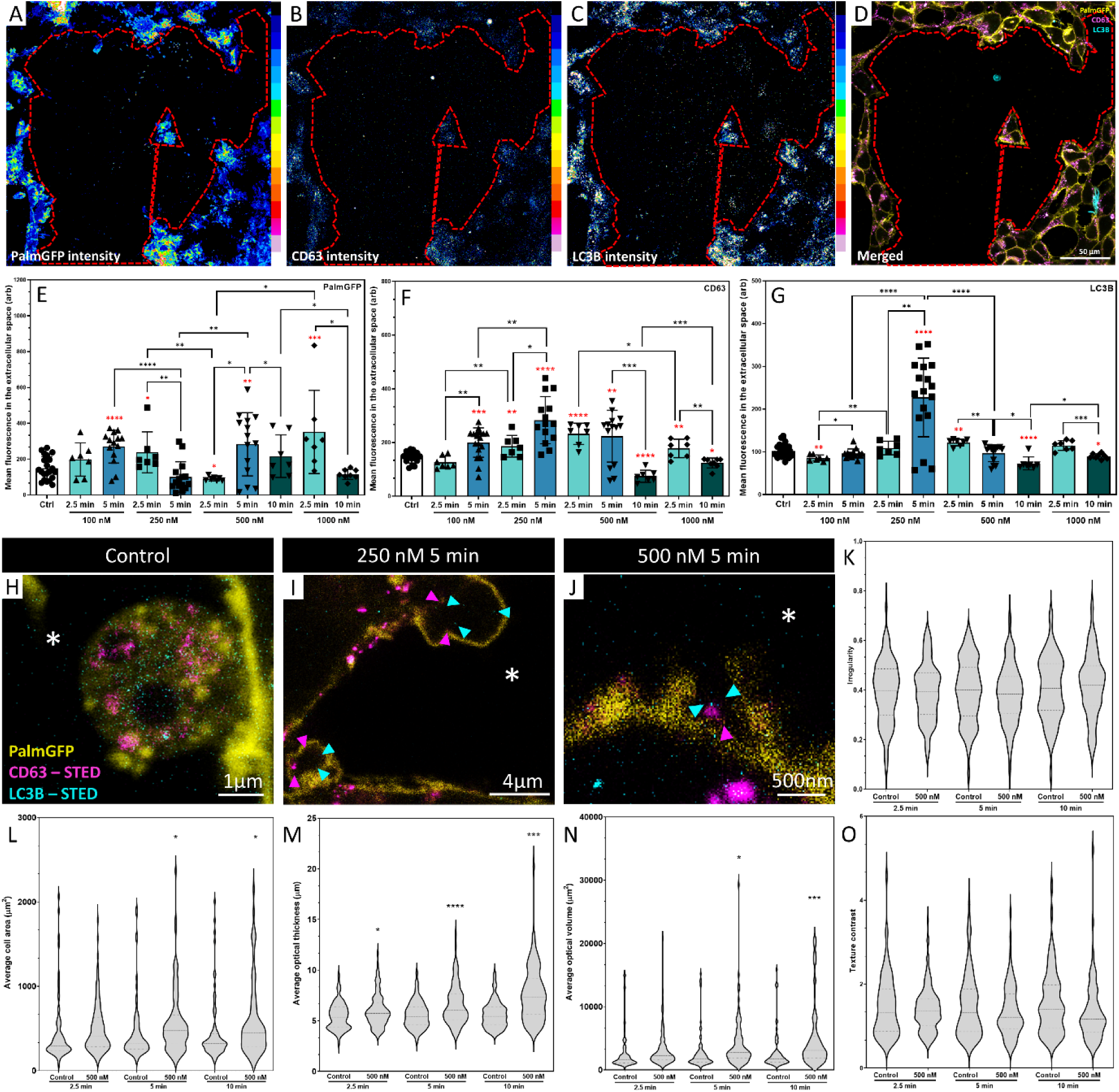
Characterization of Ca²⁺ ionophore-induced EV secretion. The mechanism and the quantity of Ca²⁺ ionophore-induced EV secretion were evaluated in HEK293T-PalmGFP cells using immunocytochemistry combined with quantitative confocal microscopy (A-G) and STED super-resolution microscopy (H-J). Mean fluorescence intensities of PalmGFP (A,E), CD63 (B,F) and LC3B (C,G) were determined in the extracellular environment. The coverslip surface (where EVs were trapped within the gelatin-fibronectin coating) is shown in representative intensity images (A-C) and a merged composite image of the cells in panel D. The extracellular space, indicated by a red dashed line (A-D) was selected based on the location of the HEK293T-PalmGFP cells (D). The analysis revealed that the concentrations of GFP-positive (plasma membrane-derived) EVs fluctuated around control levels, with some significant differences (E), whereas CD63-positive EVs followed a time- and concentration-dependent dynamics. After 2.5 min of treatment, CD63-positive EV concentrations gradually increased and saturated at 500 nM A23187. A similar trend was observed at 5 min. At 100 and 250 nM A23187 treatments, CD63-dependent fluorescence increased over time, while at 500 and 1000 nM, the opposite effect occurred. After 10 min, the signal significantly dropped (F). LC3B-positive EVs showed a pattern similar to PalmGFP-positive EVs with an additional peak at 250 nM and 5 min. At 2.5 min, a slight concentration-dependent increase was observed with saturation at 500 nM A23187 (G). STED microscopy provided evidence that at least a subset of released MV-lEVs were amphiectosomes containing CD63- and LC3B-positive intraluminal vesicles (H,I). Following 500 nM A23187 treatments for 5 min, intact amphiectosomes were not observed; however exocytosis of amphisomes were detected (J). White asterisks indicate the extracellular environments during the STED analysis. CD63 (magenta) and LC3B (cyan) were super-resolution channels while GFP (yellow) was a confocal channel. Cellular morphology and integrity were assessed during ionophore treatments of HEK293 cells using holographic microscopy. Cellular integrity (K) and texture contrast (O) remained unchanged, while average cell area (L), optical thickness (M) and optical volume (N) increased in a time-dependent manner based on quantification of 80-129 cells. Error bars represent mean ± SD. Statistical significance was determined using Student’s t-test and one-way ANOVA (*p < 0.05, **p < 0.01, ***p < 0.001, ****p < 0.0001). Red asterisks indicate statistical differences between control and treated samples.

STED imaging confirmed that at least a subset of released MV-lEVs were amphiectosomes containing CD63- and LC3B-positive intraluminal vesicles (Figure 3H,I). Intact amphiectosomes were not found after 5 min at 500 nM, although evidence of amphisome exocytosis was observed (Figure 3J).

Cellular morphology and integrity were assessed during ionophore treatment using holographic microscopy. Morphometric analysis indicated that cellular irregularity and texture contrast remained unchanged during the Ca^2+^ ionophore treatments (Figure 3K,O), while average cell area, average optical thickness and average optical volume increased significantly in a time-dependent manner (Figure 3L-N).

The effect of A23187 treatments on HEK293 cells’ viability was followed by two different assays (resazurin assay and CellTiter-Glo). Based on the resazurin assay, the treatments did not significantly change the viability (Supplementarty Figure 3A), while in CellTiter-Glo assay at 1000 nM A23187 concentration (10 min) significant drop of the ATP level was recorded (Supplementarty Figure 3B), indicating that the treatment affects the cells’ metabolic activity, but within 3h the cells were able to recover.

### Modulation of two distinct sEV release pathways

To investigate the contribution of the two distinct sEV release pathways, we combined ultrastructural analysis with targeted gene silencing and chemical modulation of either the “torn bag mechanism” or MVE exocytosis. Under steady-state conditions, TEM revealed that the predominant mechanism of sEV release was the “torn bag” pathway (Figure 4A,B), whereas MVE exocytosis was not detected. Interestingly, starvation of cells in serum-free cell culture medium (a condition commonly used in the EV field [2]), elicited a dual response: while the “torn bag mechanism” persisted (Figure 4C), MVEs accumulated in the cytoplasm (Figure 4E) and fused with the plasma membrane, indicating activation of the exocytosis pathway (Figure 4D, F).

**Figure 4.**
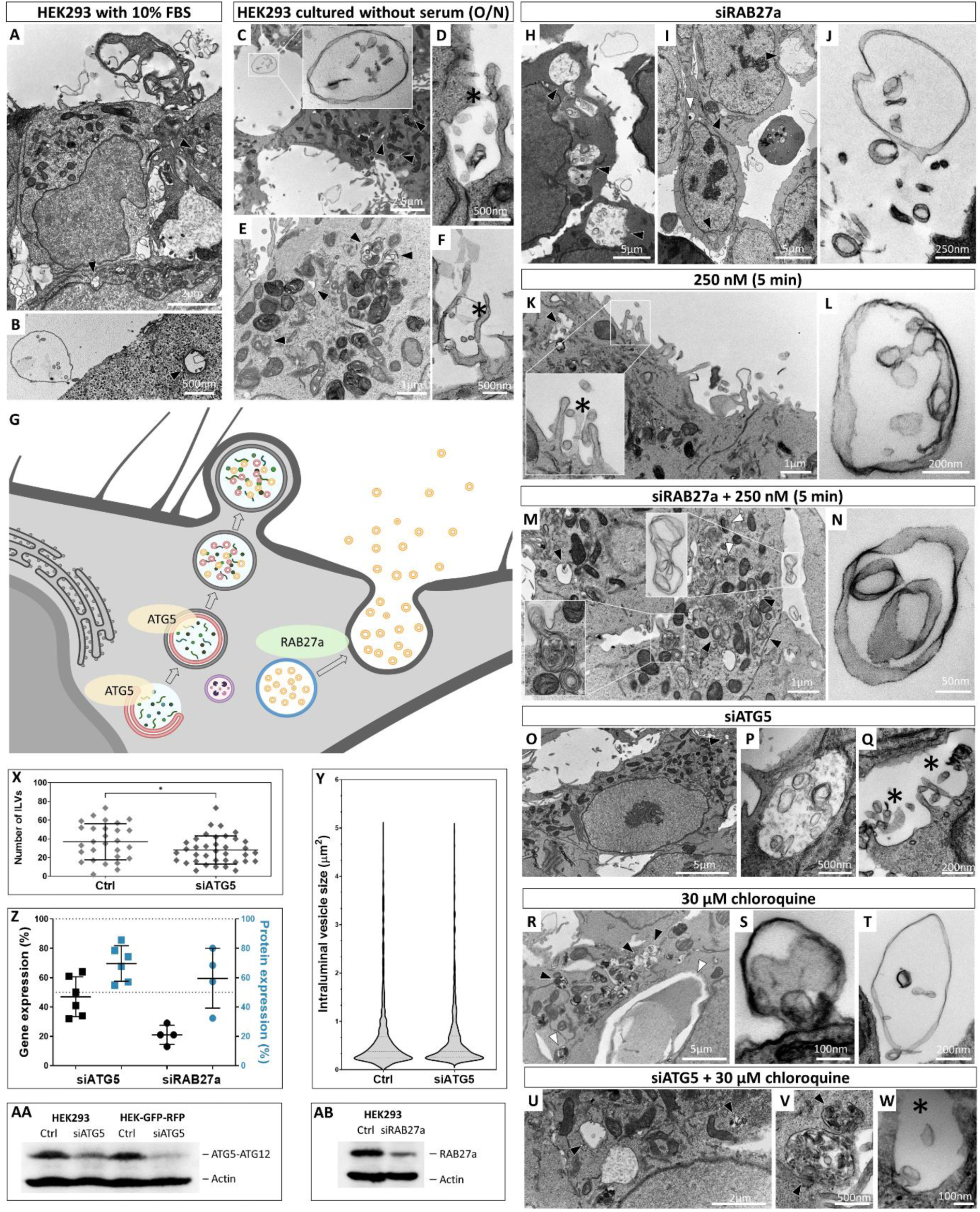
Modulation of the two distinct sEV release pathways. Serum starvation and ATG5 or RAB27a silencing were performed. Ultrastructural changes were analysed by transmission electron microscopy (A-F,H-W), while silencing efficiency was assessed by confocal microscopy in HEK293T-PalmGFP-LC3RFP cells (X,Y), RT-qPCR (Z) and western blot analysis (Z-AB). Under control conditions (A,B), the predominant sEV release mechanism was the “torn bag” pathway. In the extracellular space, intact MV-lEVs (B) and empty membrane balloons (A) were observed. Serum starvation (C-F) induced MVE exocytosis (D,F) parallel to the “torn bag” mechanism (C). MVEs accumulated in the cytoplasm (C,E) during serum starvation. The gene silencing strategy is summarized in panel G. ATG5 regulates phagophore membrane formation, whereas RAB27a is a key protein in exocytosis-based sEV secretion. Modulating the expression of these proteins enabled us to influence the two main sEV release pathways: “torn bag mechanism” and exocytosis. When RAB27a was silenced, sEV secretion occurred predominantly via the “torn bag mechanism” (H-J) while MVE exocytosis was not detected, even under 250 nM A23187 treatment (M,N). In contrast, without RAB27a silencing, we observed both the “torn bag mechanism” and the MVE exocytosis during 250 nM A23187 treatment (K,L). Silencing ATG5 abolished autophagosome formation (O), increased the abundance of MVE (P) and inducing MVE exocytosis (Q). The “torn bag mechanism” was absent and could not be reactivated even by chloroquine treatment (U-W). When chloroquine was applied alone, autophagosome formation and the “torn bag mechanism” were both activated (R-T). MVEs are indicated by black arrowheads, autophagosome by white arrowheads, MVE exocytosis by black asterisks. The effect of ATG5 silencing on autophagosome formation was confirmed by confocal microscopy using HEK293T-PalmGFP-LC3RFP cells (X,Y and Supplementary Figure 4). Gene and protein expression changes were quantified by RT-qPCR (Z) and western blot analysis (Z-AB).

To further explore the relationship between the “torn bag mechanism” and MVE exocytosis, we silenced RAB27a [27], a key regulator of MVE docking and fusion, and ATG5, an essential component of autophagosome biogenesis through phagophore membrane formation [28] (Figure 4G). RAB27a knockdown abolished MVE exocytosis (Figure 4H, I), even under calcium ionophore (A23187) stimulation (Figure 4M), forcing cells to rely exclusively on the “torn bag mechanism” for sEV release (Figure 4J, N). In contrast, in the absence of RAB27a silencing, treatment with 250 nM A23187 for 5 min induced both the “torn bag” pathway (Figure 4L) and MVE exocytosis (Figure 4K).

ATG5 silencing profoundly altered intracellular vesicular trafficking: the autophagosome formation was abolished (Figure 4O), while MVE abundance increased (Figure 4P) and exocytosis became the dominant release route (Figure 4Q). Notably, the “torn bag mechanism” was completely absent in ATG5-deficient cells and could not be induced by chloroquine treatment (Figure 4U-W), which otherwise strongly promoted both autophagosome accumulation (Figure 4R) and the “torn bag” pathway when applied alone (Figure 4S,T).

The efficiency of gene silencing and its impact on autophagy were validated by confocal microscopy using HEK293T-PalmGFP-LC3RFP cells, which confirmed a significant reduction in the number of LC3-positive particles in the cytoplasm upon ATG5 knockdown (Figure 4X), while the size distribution of LC3-positive structures remained unchanged (Figure 4Y). These findings were further supported by RT-qPCR and western blot analyses (Figure 4Z-AB), which demonstrated downregulation of ATG5 and RAB27a at both transcript and protein levels. The results suggest that sEV secretion in HEK293T cells is governed by at least two mechanistically distinct pathways.

## Discussion

Recent advances have revealed that EV biogenesis is substantially more complex than previously expected [3]. In our earlier work, we identified MV-lEVs across multiple cell lines and in murine kidney and liver, originating from amphisomes and released as amphiectosomes. These structures spontaneously rupture, and discharge intraluminal vesicles into the extracellular space via the “torn bag mechanism,” a process that likely integrates endosomal and autophagic membrane systems. Notably, under homeostatic conditions, we found no ultrastructural evidence for the conventional sEV release through MVE exocytosis [8]. The present study aimed to understand better why MVE-derived exocytosis was not observed in our previous TEM analyses and to gain further insights into the mechanisms and regulatory pathways underlying both the amphiectosome-mediated “torn bag mechanism” and MVE-driven exocytosis.

Our findings support here that sEV secretion in mammalian cells is controlled by two distinct pathways: the autophagy-dependent amphiectosome release via the “torn bag mechanism” and the classical exocytosis-dependent secretion of MVEs. These pathways may be differentially activated depending on cellular physiological conditions.

In agreement with our previous findings [8] under steady-state conditions, TEM consistently showed amphiectosome-mediated sEV release across all tested cell lines [8] and mouse tissues (Figure 1A-O, Figure 4A,B), whereas MVE exocytosis was undetectable. This suggests that the “torn bag mechanism” represents the “default” steady-state sEV secretion route in non-stressed cells. In contrast, stress conditions (including Ca²⁺ ionophore treatment and serum starvation) triggered a shift toward MVE exocytosis. Calcium influx is known to facilitate membrane repair and vesicle fusion via ANX6 [15], and our data support this model, showing rapid induction of MVE exocytosis upon A23187 treatment (Figure 2N,P-S,U-W). Based on these TEM images, we could not confirm the exclusive MVB origin of the released particles, as various EV-like structures and most probably lysosomal degraded material were present in the extracellular environment. These observations also suggest that any membrane-enclosed intracellular particle may have a membrane-protective function. Notably, the transition from a “torn bag mechanism” to an exocytosis-based sEV secretion was followed by a concentration- and time-dependent increase in extracellular particle release, as quantified by nanoparticle tracking analysis (Figure 2X-Z) and high-resolution flow cytometry (Figure 2AA). The discrepancy between scatter and fluorescence modes also suggests that a substantial fraction of released particles may lack a membrane, emphasizing that not just MVBs, but other MVEs such as lysosomes [29], autophagosomes [30] and amphisomes [9] may be involved in the exocytotic process.

Functional studies by gene silencing further clarified the molecular requirements of these pathways. ATG5 knockdown abolished amphiectosome formation through limiting autophagy [28] and the “torn bag mechanism” (Figure 4O-Q,U-W), while promoting MVE accumulation and exocytosis (Figure 4P,Q). This indicates that autophagy is needed for the biogenesis of amphiectosomes, but it is not essential for MVE exocytosis. In an opposite way, RAB27a silencing, as it was expected [27, 31], selectively blocked MVE exocytosis without affecting amphiectosome release and the “torn bag mechanism”, even under Ca²⁺ ionophore stimulation (Figure 4H-N). These results establish that the two pathways are separated and differentially regulated (Figure 5).

**Figure 5.**
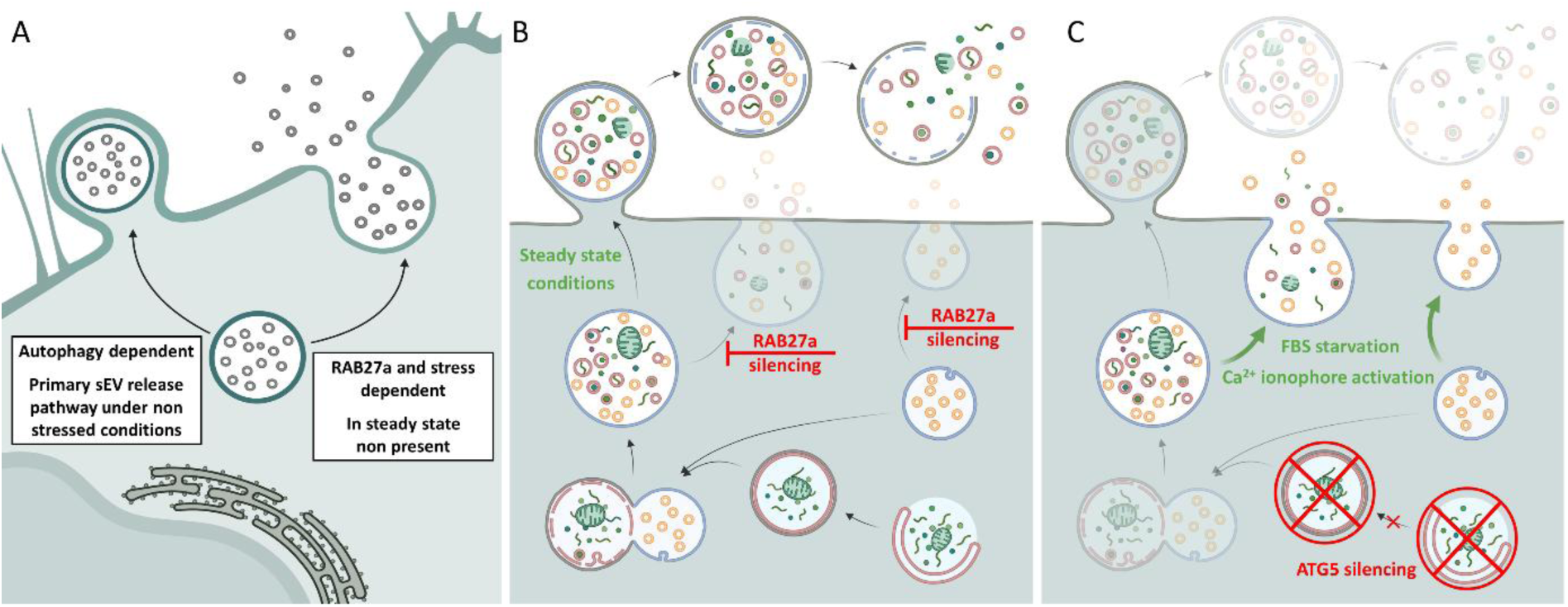
Summary of the two sEV release mechanisms. Amphiectosome release with the “torn bag mechanism” and MVE exocytosis represent two distinct sEV secretion pathways. The first pathway is autophagy dependent and is the general and the constitutive route under steady-state conditions. The second pathway appears to be more specific and is particularly activated under various stress conditions (A). RAB27a does not appear to affect either amphisome release or “torn bag”-mediated sEV secretion, but it inhibits MVE exocytosis. (B). When autophagy is inhibited, amphiectosome release is blocked and sEVs are secreted exclusively through exocytosis of MVE (C).

Our study challenges the prevailing assumption that exosome release via MVE exocytosis is the dominant route of sEV secretion under physiological conditions [11, 32–36]. Instead, we propose a model in which amphiectosome-mediated release is the primary mechanism in homeostasis, while MVE exocytosis serves as a stress-responsive pathway.

The discovery of MV-lEVs and amphiectosomes was only possible through investigations under steady-state conditions combined with *in situ* fixation. This is critical because conventional EV research and isolation protocols often rely on serum-free media [2], stress-inducing conditions [18, 37] or artificial stimulation of EV secretion [15, 38, 39], all of which can alter the origin and molecular composition of EVs. Under such non-steady-state conditions, MV-lEVs may undergo degradation, or their secretion may be replaced by MVE exocytosis. Consequently, MV-lEV secretion and sEV release via the “torn bag mechanism” can easily be overlooked. Studying MV-lEVs and amphiectosomes is challenging due to the fragile limiting membrane. Mechanical stress during pipetting or centrifugation can rupture these membranes, leaving sEVs as the predominant population in most preparations [8]. Based on the data presented here, we hypothesize that the previously reported “*en bloc*-released MVB-like EV clusters” [7], the large EVs containing ENPL [18], and the GFP-positive large EVs secreted following nanoinjection [23] likely represent amphiectosomes as well.

This paradigm shift may have significant implications for interpreting EV-based biomarkers and for designing therapeutic strategies that aim to modulate EV secretion.

Future work should address the functional consequences of these distinct EV populations, including their cargo composition and roles in intercellular communication under steady-state and stress conditions. Additionally, elucidating the signalling pathways that govern the switch between these mechanisms will be critical for understanding EV biology in health and disease.

## Acknowledgements

The authors gratefully acknowledge Anikó Gál and Györgyné Vidra for their valuable advice, as well as Soma Godó and Unicam Ltd. for making it possible to use the FluoView FV4000 confocal microscopy system. Éva Giessler, CEO of Bioxol Ltd., made the BioxML dye family available to us.

## Data Availability Statement

Data available on request from the authors.

