## Supplementary file for "Stress-induced switch in small extracellular vesicle secretion: from constitutive “torn bag mechanism” to exocytosis"

### Supplementary Figures

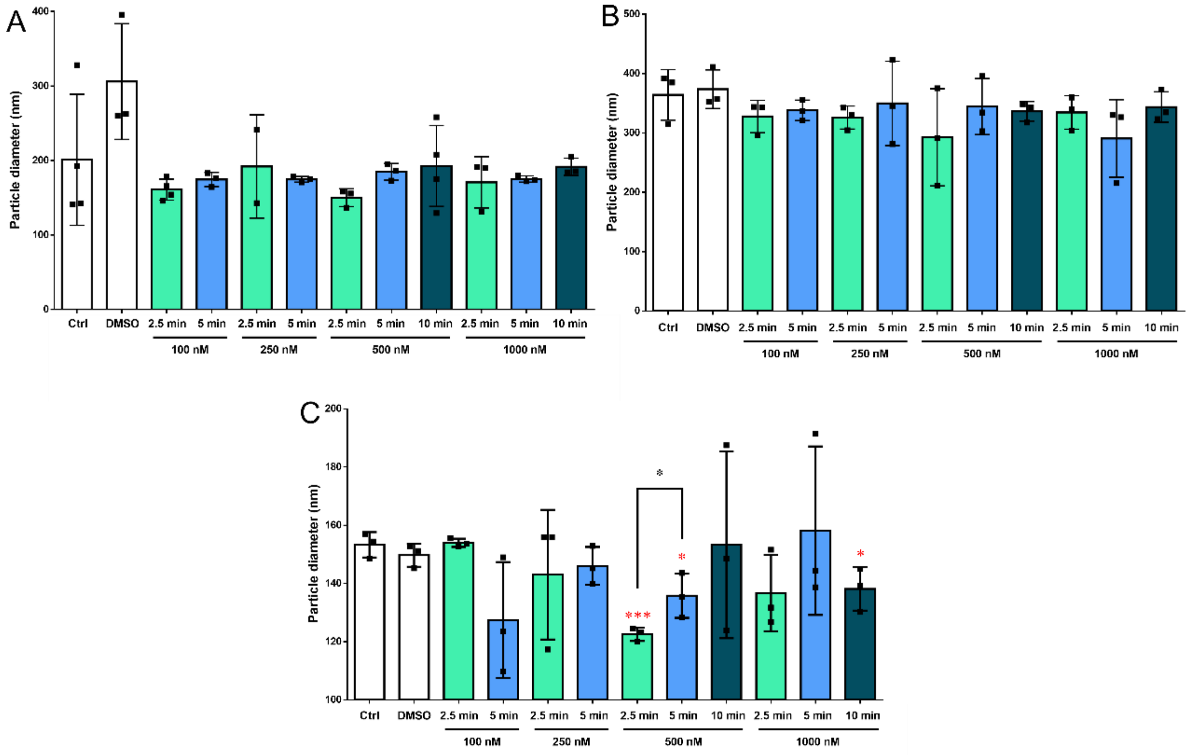


**Supplementary Figure 1. $Ca^{2+}$ ionophore treatment effects on the Median (X50) size of particles in supernatant.** Cell free supernatants were investigated with NTA in fluorescent mode (A), with settings optimised for large EVs (B) and small EVs (C). Bar charts represent mean ± SD of 3-4 independent experiments. Statistical significance was determined using Student's t-test and one-way ANOVA (*p < 0.05, ***p < 0.001). Red asterisks indicate statistical differences between control and treated samples.

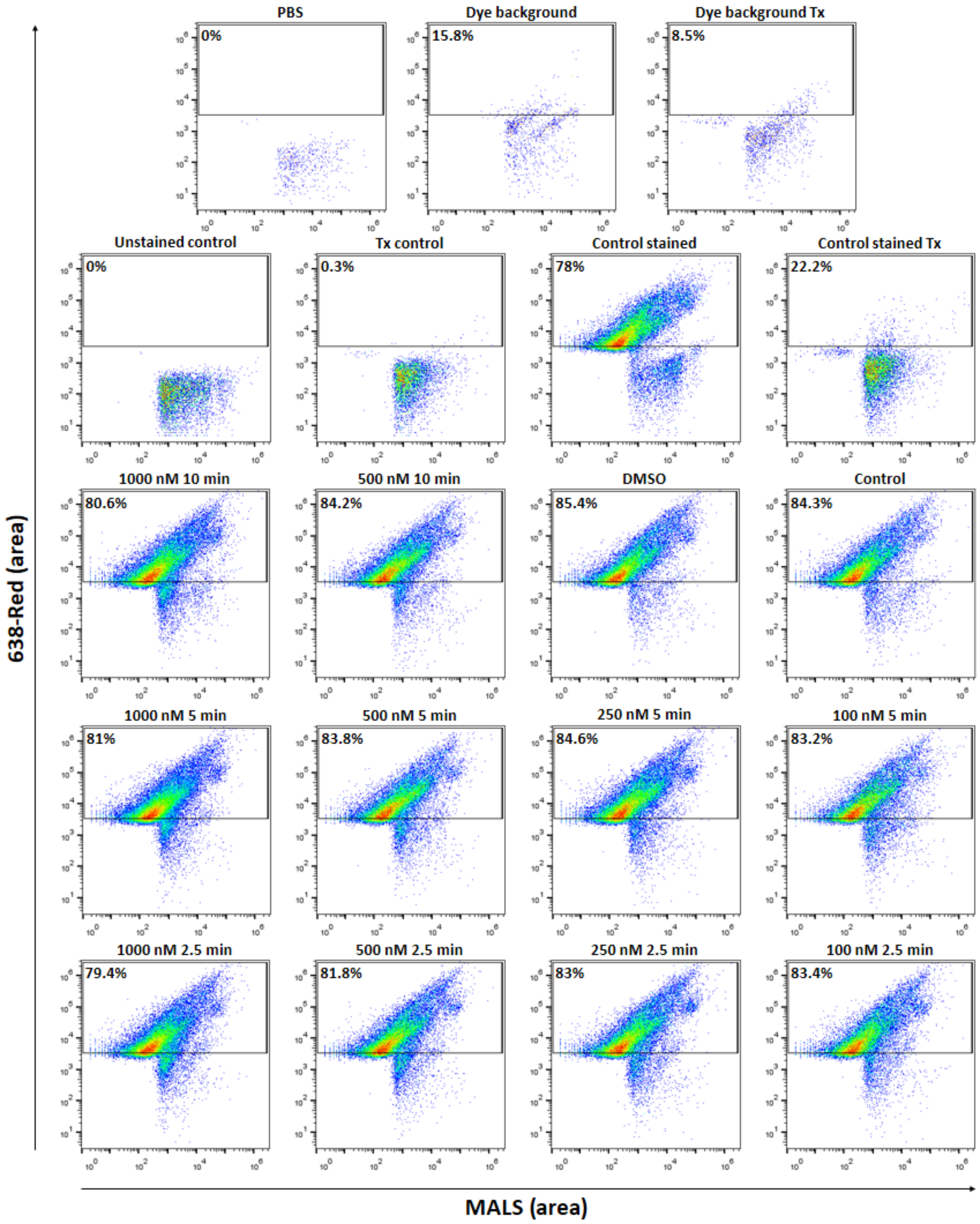


**Supplementary Figure 2. Gating strategy of EVs detection by high-resolution flow cytometry.** HEK293 cells were treated with A23187 $Ca^{2+}$ ionophore. The secreted particles in the cell-free supernatant were detected by high-resolution flow cytometry. In the separated supernatants the lipophilic dye labelled and Triton-X 100 sensitive particles were identified as EVs. Ratios of identified EVs in the total supernatant are indicated. Threshold 14 for MALS and 50 for 638-Red was applied to reduce noise. Tx (Triton-X 100)

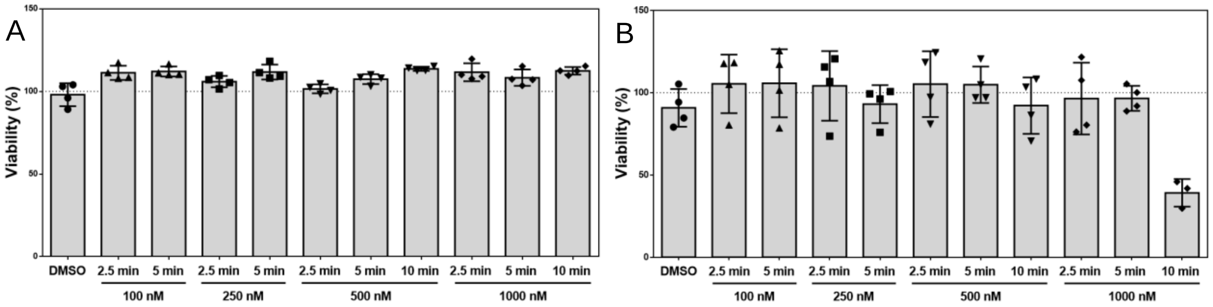


**Supplementary Figure 3. Effect of A23187 $Ca^{2+}$ ionophore treatment on cell viability.** Metabolical activity measured with resazurin assay (A) and ATP based CellTiter-Glo (B).

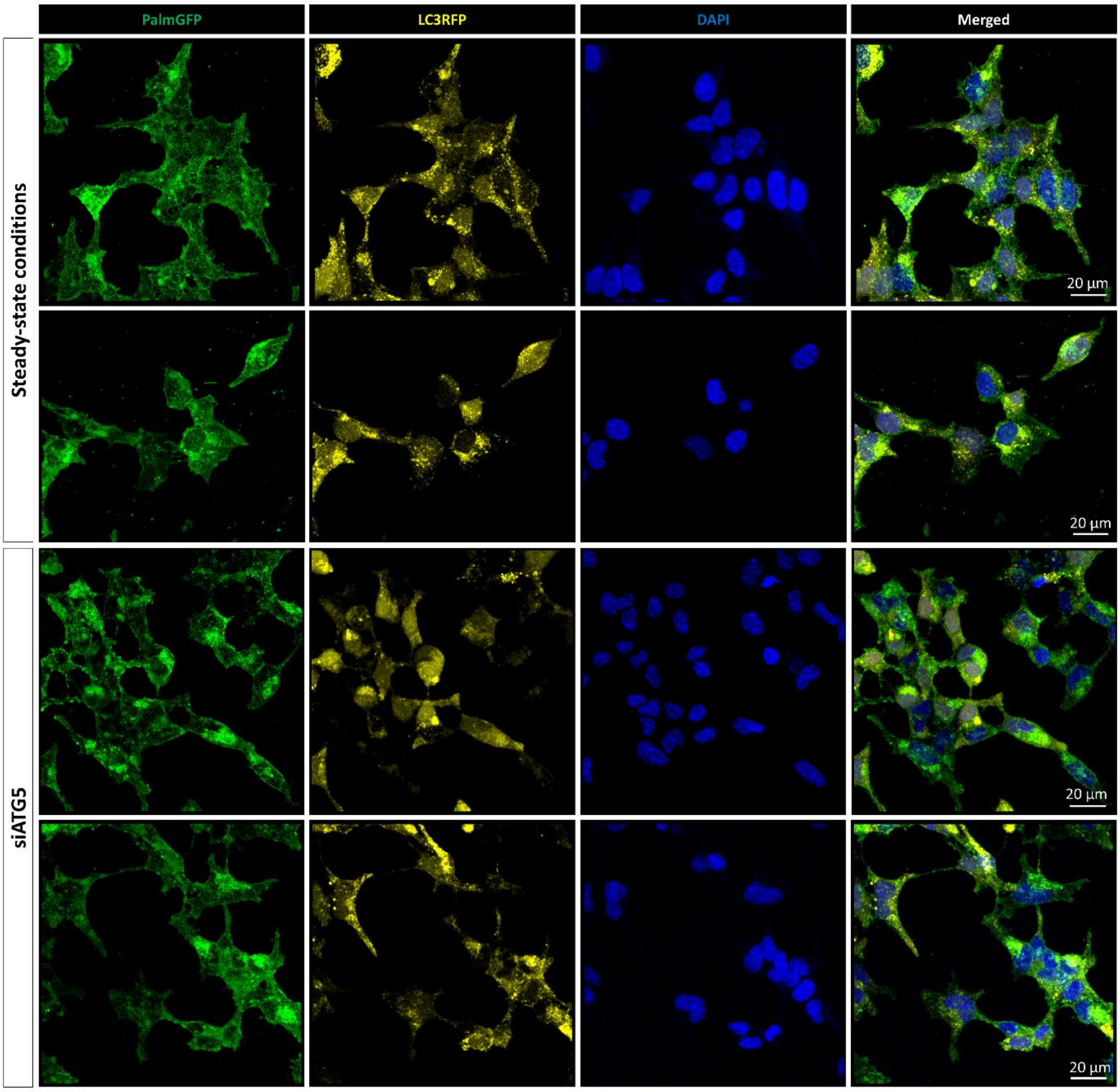


**Supplementary Figure 4. Confocal images of steady-state and ATG5-silenced HEK293T-PalmGFP-LC3RFP cells.**

### Supplementary Tables

#### Supplementary Table 1. Parameters of NTA measurements.

| | lEV | sEV | Fluorescent |
|---|---|---|---|
| **Sensitivity** | 60 | 85 | 85 |
| **Shutter** | 100 | 100 | 100 |
| **FrameRate** | 7.5 | 30 | 7.5 |
| **Positions** | 11 | 11 | 11 |
| **Cycles** | 2 | 2 | 2 |
| **MinBrightness** | 20 | 20 | 20 |
| **MinSize** | 5 | 5 | 5 |
| **MaxSize (nm)** | 1000 | 1000 | 1000 |
| **Temperature (°C)** | 25 | 25 | 25 |

**Supplementary Table 2. Configurations of high-resolution flow cytometry measurements.**

| Laser settings | | |
|---|---|---|
| | PMT | Treshold |
| LALS | 345 | |
| SALS | 350 | |
| MALS | 270 | 14 |
| 638 Red | 500 | 50 |
| Green | 525 | |
| **Flow settings** | | |
| Measuring time | 150 sec | |
| Measuring speed | 1.5 µL / min | |
| Pressure | 150 mBar | |
| **EV identification** | | |
| Lipophilic dye | 20 nM | |
| TritonX-100 | 0.1 % | |
